# AnnotateMissense: a genome-wide annotation and benchmarking framework for missense pathogenicity prediction

**DOI:** 10.64898/2026.05.03.722489

**Authors:** Muhammad Muneeb, David B. Ascher

## Abstract

**Motivation:** Missense variant interpretation remains challenging because pathogenicity depends on heterogeneous evidence, including population frequency, evolutionary conservation, transcript context, amino acid substitution severity, prior pathogenicity predictors and protein-language-model-derived features. Although these resources are individually useful, there remains a need for reproducible workflows that integrate them at genome-wide scale, benchmark their contribution and provide accessible outputs for downstream research.

**Results:** We present **AnnotateMissense**, a scalable annotation, benchmarking and genome-wide prediction framework for missense variant interpretation. AnnotateMissense integrates chromosome-wise hg38 missense variants derived from dbNSFP v5.1 with ANNOVAR-based gene-, region- and filter-based annotations, dbNSFP transcript and protein descriptors, AlphaMissense scores, ESM-derived features, conservation metrics, population-frequency variables, established pathogenicity predictors and engineered amino acid/codon-context features. Using 132,714 ClinVar-labelled missense variants, we benchmarked machine-learning and deep-learning models under controlled feature configurations. The full 303-feature benchmark set achieved the strongest performance with XGBoost, reaching mean Matthews correlation coefficient (MCC) = 0.9411 and ROC-AUC = 0.9950 across stratified five-fold cross-validation. Restricted naive and location-oriented feature sets achieved substantially lower best MCC values of 0.4989 and 0.5113, respectively. Circularity-controlled ablations showed that removing prior-predictor, population-frequency and clinically overlapping evidence reduced performance, whereas excluding AlphaMissense and ESM-derived features alone had minimal effect. Temporal ClinVar validation on newly observed pathogenic/benign variants achieved MCC = 0.7613, accuracy = 0.8798 and F1-score = 0.8750. The final genome-wide model was applied to 90,643,830 hg38 missense variants to generate AnnotateMissense pathogenicity scores and binary prediction labels.

**Availability and implementation:** Source code, workflow scripts and command files are available at https://github.com/MuhammadMuneeb007/CAGI7_Annotate_All_Missense. Genome-wide prediction outputs and the compressed DuckDB database are available at https://doi.org/10.5281/zenodo.19981867.

**Supplementary information:** Supplementary data are available online.

## Introduction

Missense variants are among the most clinically important classes of protein-altering genetic variation, but their interpretation remains difficult because functional impact depends on multiple and incompletely overlapping evidence layers. A variant may be rare in population databases, affect an evolutionarily conserved residue, alter amino acid physicochemical properties, occur in an important transcript or protein context, or be supported by prior clinical observations. No single evidence source is sufficient across all genes, proteins and disease settings, creating a persistent challenge for rare disease diagnosis, genome interpretation and large-scale variant prioritisation [1, 2, 3, 4, 5].

Many computational resources have been developed to prioritise missense variants, including conservation-based predictors, ensemble pathogenicity scores, genome-wide annotation frameworks and recent protein-language-model-derived resources [6, 7, 8, 9, 10, 11]. These methods provide complementary information, but they differ in score directionality, coverage, missingness, output type, training data and genome-build compatibility. This makes large-scale integration and systematic benchmarking difficult, particularly when some predictors may have been trained on ClinVar, HGMD or related clinical variant databases.

Here, we present **AnnotateMissense**, a reproducible framework for genome-wide missense annotation, feature integration, benchmarking, validation and prediction. Starting from 90,643,830 hg38 missense single-nucleotide variants derived from dbNSFP v5.1 through the CAGI7 Annotate-All-Missense challenge, AnnotateMissense integrates ANNOVAR- and dbNSFP-derived annotations, AlphaMissense scores, ESM-derived features, population-frequency variables, conservation scores, established pathogenicity predictors and engineered biological features [12, 13, 10, 11]. We benchmarked multiple machine-learning and deep-learning models using ClinVar-labelled missense variants, quantified feature-set contributions through controlled ablation analyses, evaluated prospective concordance using temporal ClinVar validation and generated genome-wide pathogenicity scores as a public research resource. AnnotateMissense is intended as a scalable annotation, benchmarking and research-prioritisation framework, not as a standalone clinical classification system.

## Materials and methods

### Genome-wide annotation and feature integration

AnnotateMissense was developed using chromosome-wise hg38 missense variant files derived from dbNSFP v5.1. These files comprised 90,643,830 missense single-nucleotide variants and included genomic, transcript and protein-level descriptors. Chromosome-level variant files were converted into ANNOVAR-compatible input format and annotated using gene-based, region-based and filter-based resources [12, 13]. The resulting outputs were merged with dbNSFP-derived transcript and protein context, AlphaMissense scores, ESM-derived missense effect predictions, population-frequency variables, conservation scores, established pathogenicity predictors and engineered amino acid/codon-context features [10, 11].

The integrated feature space included raw annotation variables, transformed numeric variables, categorical encodings, consensus-voting features, amino acid physicochemical change features, codon-composition variables, nucleotide substitution descriptors and selected interaction features. Direct ClinVar-derived fields not intended for prediction were excluded to reduce target leakage. Full annotation commands, database categories, feature construction details and retained/excluded feature summaries are provided in Supplementary Material 1, Supplementary Sections 1–3, and Supplementary Material 2, Sheets S1–S2.

### ClinVar benchmark and model evaluation

ClinVar-labelled missense variants were extracted from the chromosome-level annotation tables [4]. Pathogenic and likely pathogenic variants were assigned to the positive class, whereas benign and likely benign variants were assigned to the negative class. Variants annotated as uncertain significance, conflicting, ambiguous or unsupported were excluded from the primary supervised benchmark. The resulting benchmark contained 132,714 missense variants, comprising 76,804 pathogenic/likely pathogenic and 55,910 benign/likely benign variants.

Models were evaluated using stratified five-fold cross-validation. Feature preprocessing was fitted within each training fold and applied to the corresponding held-out test fold. XGBoost, Random Forest, FLAML AutoML, TabNet, a PyTorch deep neural network and a TensorFlow deep neural network were benchmarked across controlled feature configurations [14, 15, 16]. Matthews correlation coefficient (MCC) was used as the primary performance metric, with ROC-AUC, accuracy and related classification metrics used as secondary measures [17, 18].

Dataset numbering was fixed throughout the manuscript and supplementary material. Dataset 2 refers to the full 303-feature benchmark matrix, Dataset 3 to the 41-feature naive feature set, Dataset 4 to the 56-feature location-oriented feature set, and Datasets 5–8 to circularity-controlled ablation configurations. Dataset 5 excluded prior pathogenicity predictors and population-frequency metrics. Dataset 6 excluded features derived from tools trained on ClinVar, HGMD or overlapping clinical databases. Dataset 7 excluded AlphaMissense and ESM-derived features. Dataset 8 retained only engineered biological sequence-derived features. Full model settings, feature-configuration definitions and benchmark tables are provided in Supplementary Material 1, Supplementary Sections 3–6.

### Temporal validation and biological use cases

To assess generalisation beyond the original ClinVar-derived benchmark, temporal validation was performed using variants present in a newer ClinVar release but absent from the older annotation-derived ClinVar set used during model development. Variants were matched by chromosome, position, reference allele and alternate allele. Only strict pathogenic/benign categories were retained for evaluation, and uncertain or ambiguous categories were excluded. AnnotateMissense predictions were evaluated using the final categorical prediction label. Full temporal validation results are provided in Supplementary Material 1, Supplementary Section 10, and Supplementary Material 2, Sheet S4.

Biological utility was assessed using two use-case analyses. First, AnnotateMissense scores were evaluated for ClinVar variants of uncertain significance (VUS) stratified by gnomAD missense constraint Z-score [5]. Second, pairwise discordance analysis was performed against established pathogenicity predictors to identify variants where AnnotateMissense provided complementary classifications relative to existing methods. Detailed VUS prioritisation and discordance-analysis methods are provided in Supplementary Material 1, Supplementary Sections 11–12.

## Results and discussion

### AnnotateMissense provides a genome-wide missense annotation and prediction resource

AnnotateMissense generated an integrated annotation, benchmarking and prediction resource for 90,643,830 hg38 missense variants (Fig. 1). The workflow links chromosome-wise missense variant inputs to ANNOVAR-based annotations, dbNSFP transcript/protein descriptors, AlphaMissense scores, ESM-derived features, conservation metrics, population-frequency variables, established pathogenicity predictors and engineered biological variables. The final outputs include genome-wide AnnotateMissense pathogenicity scores, binary prediction labels, compressed prediction files and a queryable DuckDB database.

**Fig. 1.**
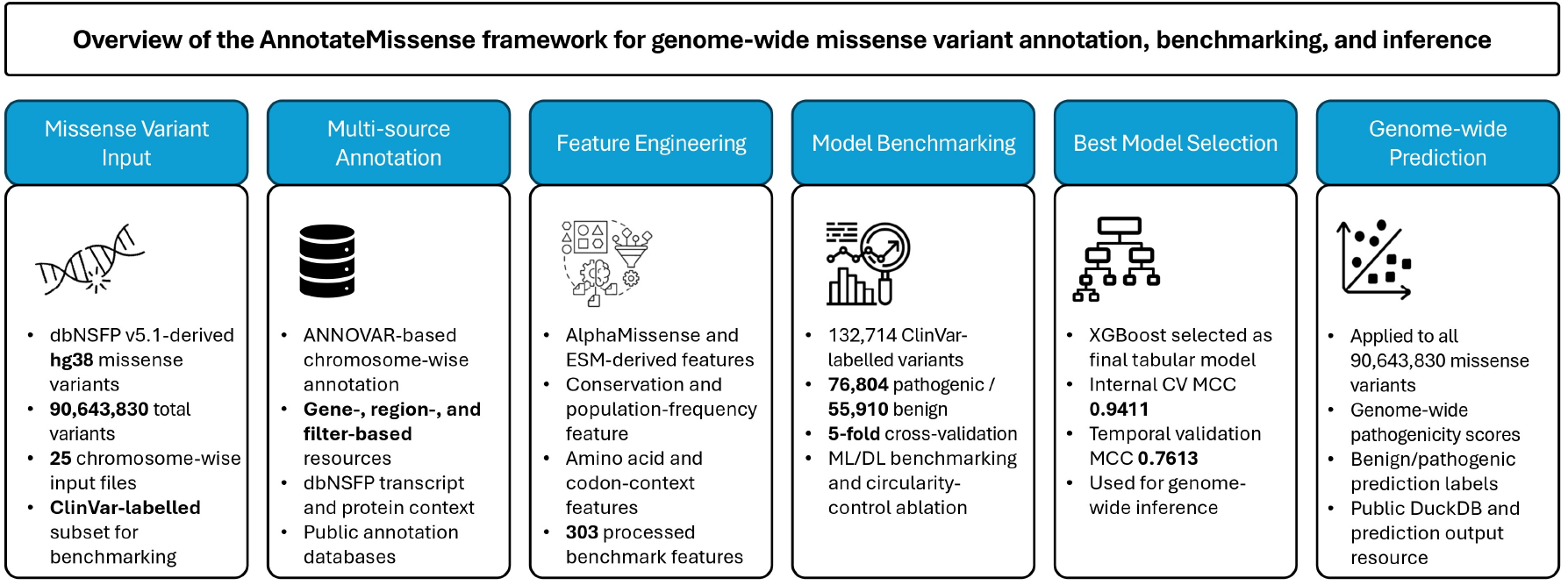
Overview of the AnnotateMissense framework for genome-wide missense variant annotation, benchmarking, validation and inference. Genome-wide hg38 missense variants derived from dbNSFP v5.1 were processed as chromosome-wise input files and annotated using ANNOVAR-based gene-, region- and filter-based resources. The resulting annotation tables were integrated with dbNSFP transcript and protein context, AlphaMissense scores, ESM-derived features, conservation metrics, population-frequency variables, prior pathogenicity predictors and engineered amino acid/codon-context features. ClinVar-labelled missense variants were used for supervised benchmarking across machine-learning and deep-learning models, with circularity-controlled ablation analyses used to assess the contribution of different evidence classes. XGBoost was selected as the final tabular model based on internal cross-validation performance, achieving an internal cross-validation MCC of 0.9411 and temporal ClinVar validation MCC of 0.7613. The final model was applied genome-wide to generate AnnotateMissense pathogenicity scores and binary prediction labels for 90,643,830 missense variants. The generated DuckDB database and prediction output files are provided as a public research resource.

### Multi-source annotation improves missense classification

Across all supervised benchmark configurations, the full multi-source feature set produced the strongest performance. On Dataset 2, XGBoost achieved the highest overall performance, with mean test MCC = 0.9411 and mean test ROC-AUC = 0.9950 across stratified five-fold cross-validation (Table 1). FLAML AutoML, Random Forest, TabNet and dense neural-network models also performed strongly on the full feature set, indicating that the integrated annotation matrix contained robust predictive signal across model families.

**Table 1.** Summary of key AnnotateMissense benchmark, ablation, temporal validation and genome-wide inference results. Full model-level results, confidence intervals, comparator analyses and machine-readable outputs are provided in Supplementary Material 1 and Supplementary Material 2.

| Analysis | Dataset / variants | Feature setting | Main result |
| --- | --- | --- | --- |
| Internal benchmark | 132,714 ClinVar variants | Dataset 2, full benchmark | XGBoost MCC = 0.9411; ROC-AUC = 0.9999 |
| Naive feature benchmark | 132,714 ClinVar variants | Dataset 3, naive features | Best MCC = 0.4989 |
| Location-oriented benchmark | 132,714 ClinVar variants | Dataset 4, location-oriented features | Best MCC = 0.5113 |
| NoPriorPredictors ablation | 132,714 ClinVar variants | Dataset 5 | Best MCC = 0.7582 |
| NoClinicalFeatures ablation | 132,714 ClinVar variants | Dataset 6 | Best MCC = 0.7326 |
| NoAlphaMissenseESM ablation | 132,714 ClinVar variants | Dataset 7 | Best MCC = 0.9409 |
| EngineeredBiologicalOnly ablation | 132,714 ClinVar variants | Dataset 8 | Best MCC = 0.5120 |
| Temporal ClinVar validation | 298,353 matched new variants | Final categorical prediction | MCC = 0.7613; F1 = 0.8750 |
| Genome-wide inference | 90,643,830 hg38 missense variants | Final genome-wide model | Public scores, labels and DuckDB resource |

Performance dropped substantially when the feature space was restricted. Dataset 3, the naive feature set, achieved a best MCC of 0.4989, while Dataset 4, the location-oriented feature set, achieved a best MCC of 0.5113. Dataset 8, which retained only engineered biological sequence-derived features, achieved a best MCC of 0.5120. These results indicate that conservation, functional annotation and local sequence-derived features alone were insufficient to recapitulate the performance of the full multi-source feature space.

Because several input features were derived from tools that may have been trained on ClinVar, HGMD or overlapping clinical variant databases, we performed circularity-controlled ablation analyses. Removing prior pathogenicity predictors and population-frequency metrics reduced performance, with Dataset 5 achieving a best MCC of 0.7582. Removing a broader set of clinically overlapping features produced a similar reduction, with Dataset 6 achieving a best MCC of 0.7326. In contrast, removing AlphaMissense and ESM-derived features alone had minimal impact, with Dataset 7 achieving a best MCC of 0.9409. These findings show that AnnotateMissense performance is driven by broad integration of heterogeneous evidence sources, while also highlighting the importance of interpreting benchmark performance in the context of predictor dependence and potential training-data overlap.

### Temporal validation and biological use cases support prioritisation utility

Temporal ClinVar validation was performed using newly observed pathogenic/benign variants not present in the older annotation-derived ClinVar set used during model development. AnnotateMissense produced categorical predictions for 298,353 newly observed pathogenic/benign variants, comprising 138,441 pathogenic and 159,912 benign variants. In this available-set temporal validation, AnnotateMissense achieved MCC = 0.7613, accuracy = 0.8798, sensitivity = 0.9070, specificity = 0.8563, precision = 0.8453 and F1-score = 0.8750. These results support prospective concordance with later ClinVar classifications, although they should be interpreted as temporal agreement with clinical variant labels rather than independent clinical validation.

AnnotateMissense was further evaluated in biological use-case analyses. Among 49,990 ClinVar missense VUS, variants in missense-intolerant genes showed higher predicted pathogenicity scores than variants in missense-tolerant genes. VUS in genes with missense constraint Z-score greater than 3 had a mean AnnotateMissense score of 0.631, compared with 0.548 for VUS in genes with missense constraint Z-score less than 0. The proportion of high-priority VUS with score greater than 0.9 followed the same gradient, supporting the biological relevance of the prioritisation scores.

Pairwise discordance analyses showed that AnnotateMissense was broadly concordant with established pathogenicity predictors while providing complementary classifications for a subset of variants. On discordant variants, AnnotateMissense showed the strongest complementary resolution against ESM1v, agreeing with ClinVar labels for 67.9% of variants where the two methods disagreed. These analyses support the use of AnnotateMissense as a research-prioritisation framework for variant triage, benchmarking and hypothesis generation.

### Framework interpretation and limitations

AnnotateMissense provides a scalable framework for genome-wide missense annotation, benchmarking, validation and prediction. The main finding is that high-performing missense classification requires integration of heterogeneous evidence sources. Models trained on the full multi-source feature set substantially outperformed restricted feature configurations based on conservation, functional annotation or local sequence-derived features alone. This supports the view that missense pathogenicity is best prioritised using combined evolutionary, biochemical, population, transcript-level and computational-predictor evidence.

The ablation analyses are central to the interpretation of the framework. Performance decreased when prior predictors, population-frequency variables and clinically overlapping evidence were removed, demonstrating that these evidence layers contribute strongly to benchmark performance. Therefore, AnnotateMissense should not be interpreted as an entirely independent clinical truth model. Rather, it is a transparent multi-source annotation and prioritisation framework that integrates existing evidence, quantifies feature-set contributions and provides scalable genome-wide inference outputs for research use.

Predictions generated by AnnotateMissense should be interpreted as prioritisation scores for research, benchmarking and variant triage, not as standalone clinical classifications. Clinical interpretation requires expert review and integration with disease context, inheritance, segregation, functional evidence and current clinical guidelines.

## Availability and implementation

The AnnotateMissense source code, workflow scripts, command files, feature-integration scripts, benchmarking code, temporal validation scripts, biological use-case analyses and genome-wide inference scripts are available at: https://github.com/MuhammadMuneeb007/CAGI7_Annotate_All_Missense

The generated genome-wide AnnotateMissense prediction resource is available from Zenodo at: https://doi.org/10.5281/zenodo.19981867

The Zenodo record includes the compressed DuckDB database `variants.duckdb.gz` and the compressed final prediction/output table `UQ_BioSig_model_Final.tsv.gz`. AnnotateMissense is provided for research prioritisation, benchmarking and variant triage, and should not be interpreted as a standalone clinical 2. classification system.

## Supplementary data

Supplementary Material 1 is available online and contains the extended annotation workflow, representative ANNOVAR commands, dataset definitions, feature configuration details, model training settings, benchmark and ablation results, temporal ClinVar validation, VUS prioritisation, discordance analysis and data/resource information.

Supplementary Material 2 is available online as a machine-readable workbook. Sheet S1 contains database information, Sheet S2 contains feature information, Sheet S3 contains prediction correlation and comparator results, and Sheet S4 contains temporal ClinVar validation results.

## Author contributions statement

M.M. wrote the first draft of the manuscript and wrote, tested, and documented the code. M.M. analysed the results. D.A. reviewed and edited the manuscript. All authors contributed to the methodology.

## Acknowledgments

D.B.A. is supported by an NHMRC Investigator Grant (GNT2041888).

## Data availability

The primary genome-wide missense variant input used in this study was derived from the CAGI7 Annotate-All-Missense challenge input based on dbNSFP v5.1. The generated AnnotateMissense genome-wide prediction outputs and compressed DuckDB database are available from Zenodo at https://doi.org/10.5281/zenodo.19981867. Source code and workflow scripts are available at https://github.com/MuhammadMuneeb007/CAGI7_Annotate_All_Missense.

